# RIG-I has a role in immunity against *Haemonchus contortus*, a gastrointestinal parasite in *Ovis aries- a novel report*

**DOI:** 10.1101/2020.09.30.320218

**Authors:** Samiddha Banerjee, Aruna Pal, Abantika Pal, Subhas Chandra Mandal, Paresh Nath Chatterjee, Jayanta Kumar Chatterjee

## Abstract

RIG-I is associated to the DExD/H box RNA Helicases. It is a Pattern Recognition Receptor (PRR), playing a crucial role in the system and is a germ line encoded host sensor to perceive Pathogen Associated Molecular Patterns or PAMPs. So far reports were available for the role of RIG-I in antiviral immunity. This is the first report we have documented the role of RIG-I in parasitic immunity. *Haemonchus contortus* is a deadly parasite affecting sheep industry which has a tremendous economic importance and the parasite reported to be prevalent in the hot and humid agroclimatic region. We had characterized RIG-I gene in sheep (Ovis aries) and identified the important domains or binding site with *Haemonchus contortus* through *in silico* studies. Differential mRNA expression analysis revealed upregulation of RIG-I gene in abomassum of infected sheep compared to that of healthy sheep, further confirming the findings. Thus it is evident that in infected sheep, expression of RIG-I is triggered for binding to more pathogen (*Haemonchus contortus*). Genetic similar studies with human and other livestock species were conducted to reveal that sheep may be efficiently used a model organism for studying the role of RIG-I in antiparasitic immunity of human.

## 1. Introduction

The innate immune system acting as the primary step of defense against infectious agents recognizes the Pattern Recognition Receptors (PRRs). They play a crucial role in the system and are germ line decoding host sensors to decipher Pathogen Associated Molecular Patterns or PAMPs and are expressed by innate immune cells like dendritic cells, macrophages, monocytes, neutrophils etc. RIG-1 (Retinoic acid inducible gene-I) like receptors that comes under the PRR superfamily detects viral nucleic acid in the cytosol^**1**^. RIG-I is associated to the DExD/H box RNA Helicases and is one of the members of RIG-I like Helicases, whereas the other two are MDA5 and LGP2. RIG-I is intimately in association with the Dicer family of helicases of RNAi pathway.

RIG-I preferably recognizes short RNA sequences marked with 5’-triphosphate group (5’ppp) and blunt end of short double stranded or single stranded RNA (ds RNA) or single stranded RNA (ss RNA) hairpins ^**2**^. It binds mostly with negative strand viruses, e.g. influenza A viruses and in some positive and double strand RNA viruses also ^**3**^. RIG-I is crucial for signaling by Influenza A, influenza B, Human respiratory syncytial virus, paramyxovirus, Japanese encephalitis virus and West Nile virus.

The molecular kinetics of RIG-I contains two N-terminal CARD Domains and one RNA helicase domain which helps in relaying the signal to the downstream signal adaptor Mitochondrial antiviral signaling protein or MAVS which tends to drive type I IFN responses. It also induces caspase – 8 dependent apoptosis preferably in tumor cells ^**4**^. Tumor cells are highly vulnerable to RIG-I induced apoptosis while normal healthy cells are resistant to RIG-I induced apoptosis. RIG-I gene participates in TLR stimulated phagocytosis ^**5**^. When RIG-I is stimulated with TLR4, it induces its expression in macrophages and gradually depletion of RIG-I causes inhibition of TLR4 induced bacterial phagocytosis. Investigations report that RIG-I acts as a potential therapeutic target for cancer presicely melanoma. In the context of viral infection, RIG-I induces MAVS dependent inflammasome activation^**6**^. Studies have shown that RIG-I play a vital role in immune responses in association with different infectious and non infectious diseases like astherosclerosis, skin psoriasis in which the levels of Interferon gamma and RIG-I is significantly raised in the epidermis of the skin ^**7**^.RIG-I expression can be highly observed in differentiated skin and colon mucosal tissue.

Indigenous sheep (*Ovis aries*) which forms a back bone of socio economically backward farmers and landless labourers and has become their way of livelihood. Nematode infection as infection with *Haemonchus contortus* possess a major threat to sheep industry both in commercial farms or at farmers herd. Earlier we had studied the genetic resistance of indigenous sheep with respect to mitochondrial genes ^**8**^. So far informations are available for role of RIG-I on viral infection, as well as few reports of antibacterial immunity. Till date reports are not available for role of anti parasitic effect of RIG-I.

Hence, the present study reflects on the information regarding the structure, function and expression profile of RIG-I gene in indigenous sheep through molecular methods and the role of RIG-I gene in parasitic infection in healthy and diseased sheep for the first time.

## 2. Materials required and Methods

### 2.1. Animals, Collection of Samples and mRNA Isolation

#### Animals, faecal sample collection and determination of FEC

We had randomly collected 60 Garole sheep samples from ILFC farm, WBUAFS, Mohanpur campus. The samples were collected prior to routine deworming procedure, presumed to have pre existing gastro intestinal parasite since they were not dewormed for last three months for the purpose of the present study during the monsoon season. The animals were presumed to get exposed to natural infection during grazing.

Faecal samples of the sheep were examined thoroughly by salt floatation method and the Faecal Egg Count were screened for each and every sample. Based on statistical analysis, samples were collected from two groups, designated as healthy (x-bar+SD), where x-bar stands for mean and SD means Standard Deviation) and diseased (x-bar – SD), where x-bar stands for mean and SD stands for Standard Deviation). Sheep were regularly sold and slaughtered for the purpose of mutton production unit. Altogether 12 t issue sampes were screened, 6 each with low FEC (grouped as healthy) for further case study. Tissue samples were collected for abomasum, rumen, small intestine,caecum, liver and lymph node.

#### mRNA isolation

Tissues from different body organs from healthy and disease sheep were collected and subjected to total RNA isolation by Trizol method. These are primarily the organs of digestive system as abomasum, rumen, small intestine, caecum and liver to assess gut associated lymphoid tissue (GALT). Other tisues studied were lymph node, testis, urinary bladder, kidney, inguinal lymph node and heart. Abomasum tissue were collected individually from healthy sheep and diseased sheep (*Haemonchus controtus* affected). mRNA isolation was carried out aseptically by Trizol method. Each healthy and diseased abomasums tissue collected were marked and separated as; tip of abomasum part and middle of abomasum part. mRNA was withdrawn from tissue of abomasum by Trizol method and subjected to cDNA preparation^**9,10, 11**^. Each mixture of 2g of tissue and 5ml trizol was triturated, followed by chloroform treatment. Centrifugation resulted in the three phase differentiation stage from which the aqueous phase containing the mRNA was separated. It was treated by isopropanol to extract the mRNA in the form of a pellet discarding the supernatant as a result of centrifugation. Finally, the pellet under went a ethanol wash and was air dried followed by dissolving the mRNA in nuclease free water. RNA concentration and quality was estimated by Nanodrop as per standard procedure.

#### Materials required

10X buffer, dNTP and Taq DNA polymerase were purchased from Invitrogen, SYBR Green qPCR Master Mix (2X) was purchased from Thermo Fisher Scientific Inc. (PA, USA). Primers were purchased from Xcelris Labs Limited. The reagents used were of analytical grade.

### 2.2. Amalgamation,Configuration of cDNA and PCR Amplification of RIG-I gene

20 μL reaction mixture volume consisted of 5 μg of complete RNA, 40 U of Ribonuclease inhibitor, 0.5 μg of oligo dT primer (16–18 mer),1000 M of dNTP, 5 U of MuMLV reverse transcriptase in reverse transcriptase buffer and lastly 10 mM of DTT. The reaction mixture was carefully blended and incubated properly at 37°C for 1 hour. The reaction was ceased and heated at 70°C for 10 minutes and immediately chilled on ice. The probity of the cDNA checked by undergoing the Polymerase Chain Reaction. To obtain full length open reading frame (ORF) of gene sequence, specific primers pairs were designed and confirmed based on the mRNA sequences of *Bos indicus* by DNASTAR software as registered in **Table1**. The amplified products were the overlapping sequences joined to extract the full sequence.

**Table 1:**
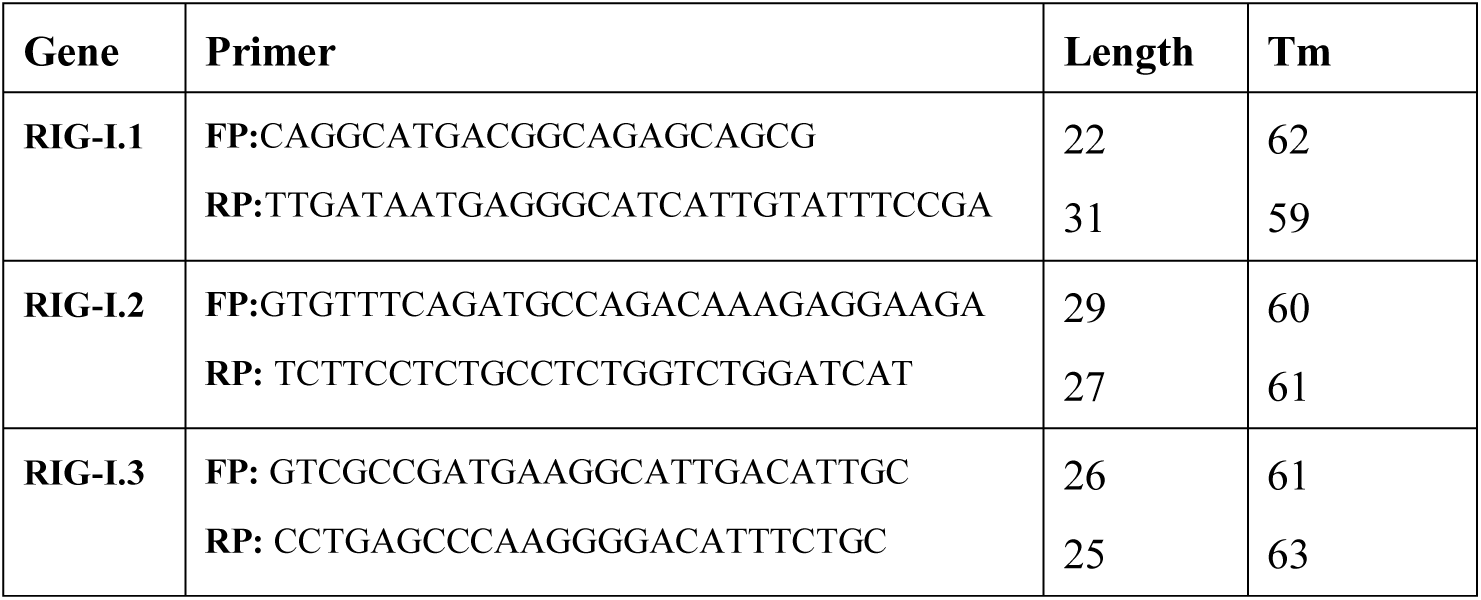
List of primers used for amplification of RIG-I gene of Garole sheep as overlapping sequences.

### 2.3. Sequence Analysis

The nucleotide sequence so extracted was scrutinized for protein translation, contigs comparisons and sequence alignments by DNASTAR Version 4.0, Inc.,USA. Novel sequence was submitted to the NCBI Genbank and accession numbers were obtained which is available in public domain (ACCESSION No. KX687005).

### 2.4. Study of Predicted ovine RIG-I peptide Using Bioinformatics Tools

The edited peptide sequence of RIG-I gene of Garole sheep was obtained by Lasergene Software, DNASTAR and then alignment of the RIG-I peptide of other rodent species and *Homo sapiens* using MAFFT^**12**^ was performed.

Prediction of signal peptide of RIG-I gene was derived by using the Signal P 3.0 Sewer-prediction results software, Technical University of Denmark. Calculation of Leucine percentage was done manually from predicted peptide sequence. Di-sulphide bonds were derived using proper and suitable software (http://bioinformatics.bc.edu/clotelab/DiANNA/) and homology searching with other species^**13**^.

Detection of Leucine-rich nuclear export signals (NES) was obtained with NetNES 1.1 Server, Technical university of Denmark. O-linked glycosylation sites analysis was carried out using NetOGlyc 3.1 server (http://www.expassy.org/), whereas N-linked glycosylation detection was done by NetNGlyc 1.0 software (http://www.expassy.org/). Protein sequence level analysis study was carried out by Blast (http://www.expasy.org./tools/blast/) for determination of leucine rich repeats (LRR), leucine zipper, N-linked glycosylation sites, detection of Leucine-rich nuclear export signals (NES), and detection of the position of GPI anchor.

α- helix and β- sheet regions were predicted using NetSurfP-Protein Surface Accessibility and Secondary Structure Predictions, Technical University of Denmark ^**14**^. Detection of Leucine-zipper was obtained through Expasy software, Technical university of Denmark. Domain linker prediction was carried out according to the software developed ^**15**^. LPS binding and LPS signalling sites are essential factor for innate immune function as pathogen recognition and binding; LPS-binding sites as well as LPS-signalling sites ^**16**^ were predicted based on homology studies with other species CD14 polypeptide.

### 2.5. Model quality assessment and three dimensional structure prediction

The templates which contain the highest sequence identity with our target template were identified by using PSI-BLAST (http://blast.ncbi.nlm.nih.gov/Blast). PHYRE2 server^**17**^ was used for homology modeling and 3D structure based on homologous template structures were detected. Subsequently, the mutant model was generated using PyMoL tool. The Swiss PDB Viewer was employed for controlling energy minimization. The 3D structures were analysed by PyMOL (http://www.pymol.org/) which is an open source molecular visualization tool.. The structural evaluation along with stereo-chemical quality assessment of predicted model were obtained by using the Structural Analysis and Verification Server (SAVES), which is an integrated server (http://nihserver.mbi.ucla.edu/SAVES/). The ProSA (Protein Structure Analysis) web server (https://prosa.services.came.sbg.ac.at/prosa) was used for clarifying and validation of protein structure. The ProSA was used for overall prospective and checking the model structural quality with potential errors and program shows a plot of its residue energies and Z-scores which determine overall quality of model ^**18**^. TM align software was used for 3 D structure alignment of IR protein for different species and RMSD prediction to generate the structural differentiation ^**19**^. The solvent accessibility surface area of the IR genes was generated by using NetSurfP server (http://www.cbs.dtu.dk/services/NetSurfP/) ^**14**^. It calculates relative surface accessibility, Z-fit score, probability for Alpha-Helix, probability for beta-strand and coil score, etc.

### 2.6. Protein-protein interaction network depiction

In order to analyze the network of RIG-I peptide, we performed the analysis with submitting FASTA sequences to STRING 9.1 ^**20**^. In STRING, the functional interlinkage was analyzed by using confidence score. Interactions with score < 0.3 are considered as low confidence, scores ranging from 0.3 to 0.7 are classified as medium confidence and scores > 0.7 yield high confidence. The functional partners were illustrated.

### 2.7. Molecular Docking

Molecular docking is a bioinformatics tool used for *in silico* analysis for the prediction of binding mode of a ligand with a protein 3D structure. Patch dock is an algorithm for molecular docking based on shape complementarities principle ^**21**^.

### 2.8. Differential mRNA expression profiling of ovine RIG-I with Real Time PCR (qRT-PCR)

#### Differential mRNA expression profile was conducted in two phases

Differential mRNA expression profiling of RIG-I gene was conducted for different organs of sheep in first phase. These are abomasum, duodenum, rumen, caecum and liver among the digestive system. Other organs are heart, kidney, urinary bladder, lymph node, testis, inguinal lymph node. RIG-I expression was observed to be important for gut associated lymphoid tissue (GALT). RIG-I was expressed mostly in the abomasum of sheep.

Consideration of the above findings, and since Haemonchus mostly infects the abomasum, in the phase, we had studied differential mRNA expression profiling of Abomasum in healthy and Haemonchus infected sheep.

All qRT-PCR reactions were conducted on ABI 7500 system. Equal amount of RNA (quantified by Qubit fluorometer, Invitrogen), wherever required, were used for cDNA preparation (Superscript III cDNA synthesis kit; Invitrogen).. Each reaction mixture consisted of 2 µl cDNA, 10 µl of single strength SYBR Green PCR Master Mix, 0.5 µl of each forward and reverse primers (10 pmol/µl) and nuclease free water for a final volume of 7 µl. Each sample was run in duplicate. Analysis of real-time PCR (qRT-PCR) was performed by delta-delta-Ct (ΔΔCt) method. The list of primers used for QPCR study have been listed below in **Table 2**:

**Table 2:**
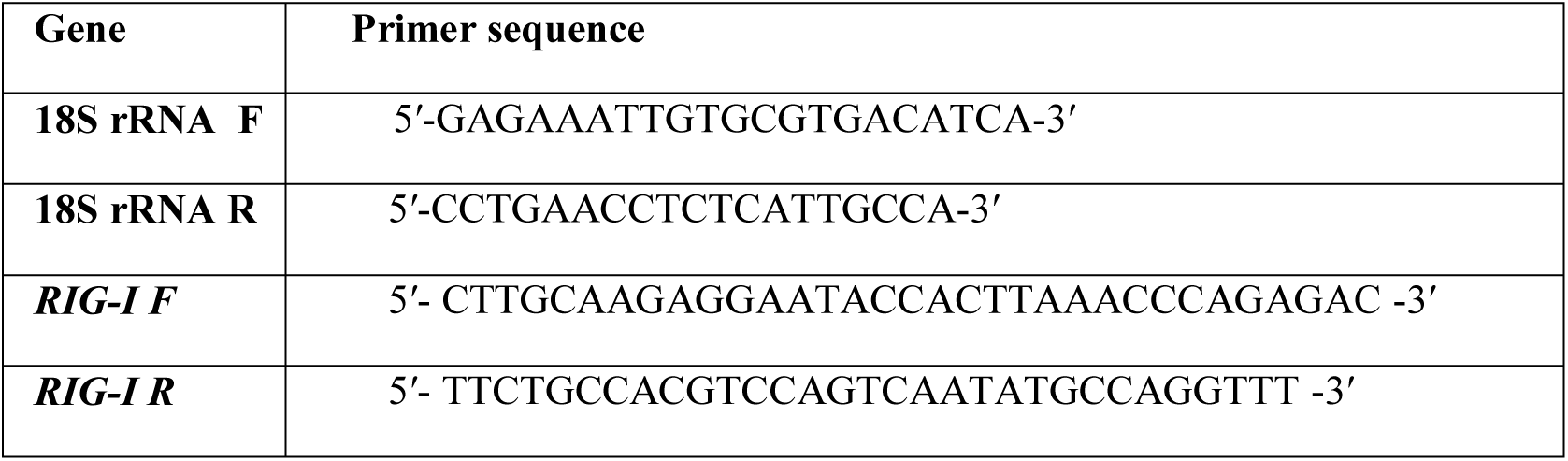
List of primers used for QPCR study.

### 2.9. Phylogenetical analysis

Nucleotides as well as derived amino acid sequences were then aligned with that of the reported sequences of different species derived from Gene Bank (http://www.ncbi.nlm.nih.gov/blast) for RIG-I Gene. Phylogenetic analysis was conducted with MAFFT software to determine the evolutionary relationship. A neighbor-joining method is employed to reconstruct phylogeny for the putat ive alignment with the software as MAFFT ^**12**^after the multiple alignment is completed.

### 2.10. Statistical Analysis

Computational descriptive statistics were done through Microsoft Excel. Analysis of variance (ANOVA). Furthermore it was used to test between groups and intervals between hours. Fischer’s restricted least significant differences criterion was used to keep the a prior type I error rate of 0.05. All statistical analysis were done out using SYSTAT 13.1 software (SYSTAT Software Inc.).

## 3. Results

### 3.1. Assessment of parasitic infestation in sheep

Screening of the faecal egg count (FEC) of the sheep (n=60) lead to study the health condition of the sheep which further prevailed us to divide them into two categories as Healthy and Diseased group. The mean FEC of the animal are given in the **Table 3**.

**Table 3:** Mean FEC of the Healthy and Diseased sheep:

| Health Status of the sheep | Mean Faecal Egg Count |
| --- | --- |
| Healthy | 50± 5.56 |
| Disease | 550± 9.45 |

### 3.2. Molecular Characterization of RIG-I gene of sheep

RIGI gene of sheep has been characterized and sequence obtained is submitted to genebank (Accession number KX687005). The 3D structure have been depicted in **Figure 1a**. The surface view of RIG-I gene has been given in **Figure 1b** which has been extracted from the PyMol software. Using different bioinformatics tools several important domains of the gene and their functions are listed in **Table 4:**

**Figure 1a:**
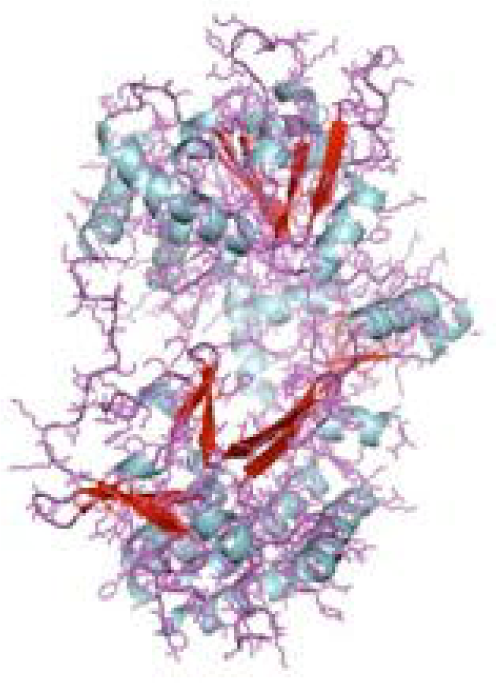
3D Structure of RIG1 gene of sheep.

**Figure 1b:**
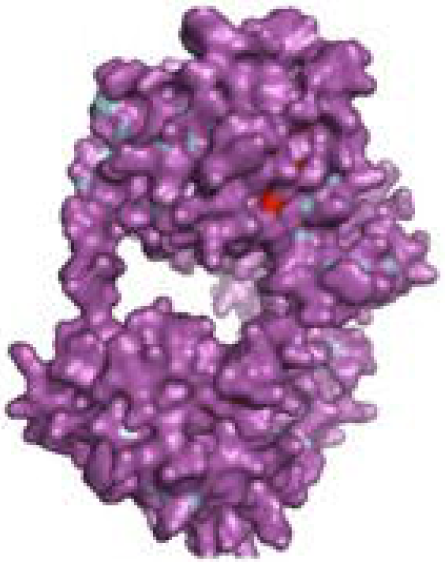
3D Structure of RIG1 gene (surface view) of sheep.

**Table 4:** List of the important domains of RIG-I of sheep assessed through *in silico* study.

|  |  |  |  |
| --- | --- | --- | --- |
| RIG-I_C-RD | C- terminal domain of RIG-I | 772-886 | Fig 2 |
| RIG-I_C | C-terminal domain of Retinoic acid-inducible gene (RIG)-I protein | 769-882 | Fig 3 |
| CARD_RIG-I_r1 | Caspase activation and recruitment domain found in RIG-I | 2-92 |  |
| CARD_2 | Caspase recruitment domain | 1-93 |  |
| RLR_C | C-terminal domain of Retinoic acid inducible gene-I | 771-882 | Fig 4 |
| RLR_C_like | C-terminal domain of Retinoic acid-inducible gene | 772-856 | Fig 5 |
| CARD_RIG-I_r2 | Caspase activation and recruitment domain found in RIG-I | 100-160 |  |
| CARD_IPS-1_RIG-I | Caspase activation and recruitment domains (CARDs) found in IPS-1 and RIG-I-like RNA helicases | 2-92 |  |
| CARD_2 | Caspase recruitment domain | 99-160 |  |
| LGP2_C | C-terminal domain of Laboratory of Genetics and Physiology 2 (LGP2) | 771-883 | Fig 6 |
| MDA5_C | C-terminal domain of Melanoma differentiation-associated protein 5 | 772-883 | Fig 7 |
| CARD_IPS-1_RIG-I | Caspase activation and recruitment domains (CARDs) found in IPS-1 and RIG-I-like RNA helicases | 100-183 |  |
| CARD_MDA5_r1 | Caspase activation and recruitment domain found in MDA5, first repeat; Caspase activation | 12-81 |  |
| MPH1 | ERCC4-related helicase [Replication, recombination and repair]; | 678-769 | Fig 8 |
| CARD_RIG-I_r2 | Caspase activation and recruitment domain found in RIG-I, second repeat | 16-93 |  |
| CARD_MDA5_r2 | Caspase activation and recruitment domain found in MDA5, second repeat | 41-87 |  |
| CARD | Caspase activation and recruitment domain: a protein-protein interaction domain | 6-87 | Fig 9 |
| PRK13766 | Hef nuclease; Provisional | 678-771 | Fig 10 |

Firstly, the RIG-I_C-RD domain (772-886), the C-terminal domain of RIG-I has been shown as the yellow portion in the **Figure 1c**, RIG-I_C(769-882), C-terminal domain of Retinoic acid-inducible gene (RIG)-I protein has been shown as a blue portion in the **Figure 1d**.

**Figure 1c:**
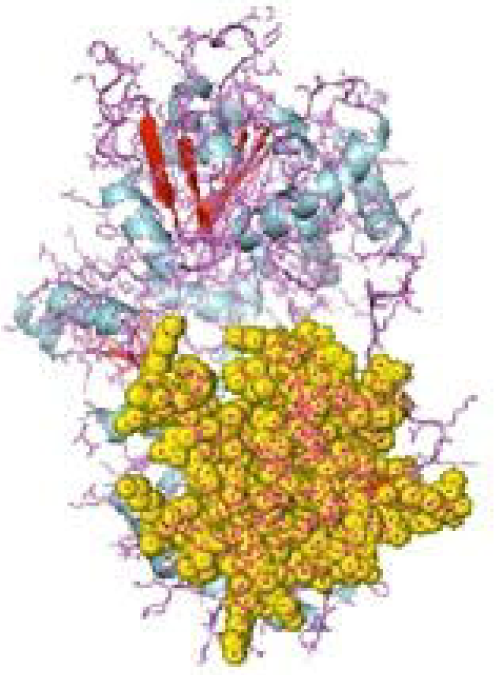
RIG-I_C-RD, the C- terminal domain of RIG-I of sheep.

**Figure 1d:**
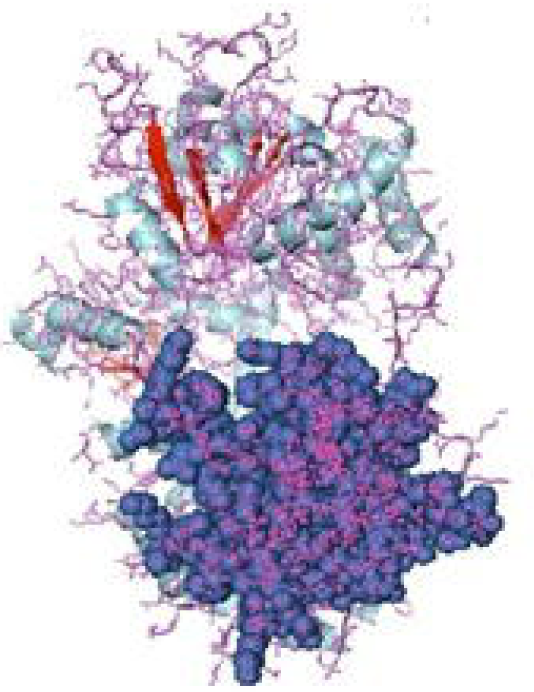
RIG-I_C, C-terminal domain of Retinoic acid-inducible gene (RIG)-I protein of sheep.

CARD_RIG-I_r1(2-92) Caspase activation and recruitment domain found in RIG-I .CARD_2 (1-93) which is also a Caspase recruitment domain is present somewhat in the similar portion.

RLR_C (771-882) is a C-terminal domain of Retinoic acid inducible gene-I depicted in **Figure 2a** as an orange colour portion.

**Figure 2a:**
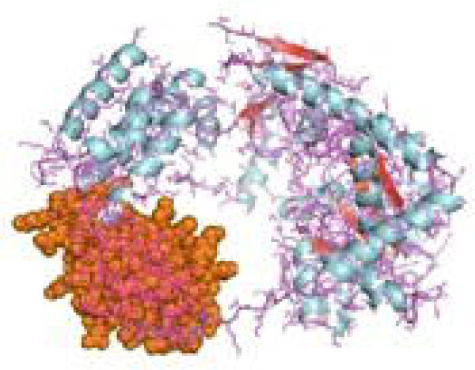
RLR_C terminal Domain of the gene of sheep.

RLR_C_like domain (772-856) C-terminal domain of Retinoic acid inducible gene-I in **Figure 2b** is depicted in the green colour part.

**Figure 2b:**
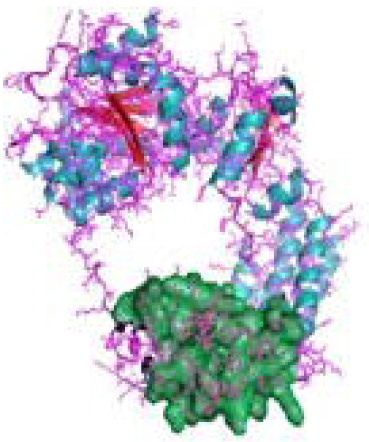
RLR_C_like domain of RIG gene of sheep.

CARD_RIG-I_r2 (100-160) is the Caspase activation and recruitment domain found in RIG-I. Here again 2-92 portion is the CARD_IPS-1_RIG-I domain i.e. Caspase activation and recruitment domains (CARDs) found in IPS-1 and RIG-I-like RNA helicases. CARD_2 (99-160) is the Caspase recruitment domain of the gene.

LGP2_C (771-883) which is in brown colour in the **Figure 2c** is the C-terminal domain of Laboratory of Genetics and Physiology 2 (LGP2). **Figure 2d** highlighted with the deep blue colour is the MDA5_C (772-883), that is the C-terminal domain of Melanoma differentiation-associated protein 5.

**Fig 2c:**
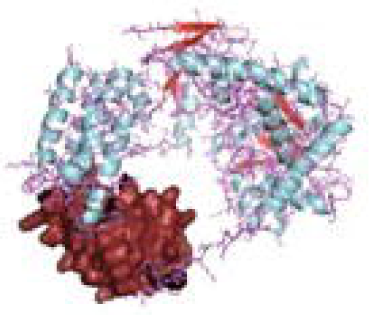
LGP2_C, C-terminal domain of Laboratory of Genetics and Physiology 2 (LGP2) of sheep.

**Fig 2d:**
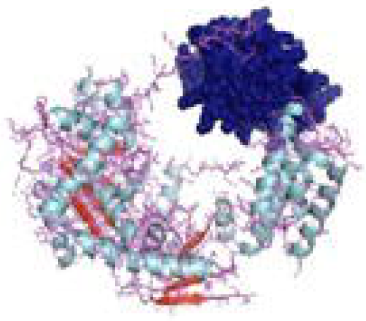
MDA5_C, C-terminal domain of Melanoma differentiation-associated protein 5 of sheep.

CARD_IPS-1_RIG-I (100-183) is the Caspase activation and recruitment domains (CARDs) found in IPS-1 and RIG-I-like RNA helicases. CARD_MDA5_r1 (12-81) represents the Caspase activation and recruitment domain found in MDA5, first repeat; Caspase activation.

MPH1(678-769) showing in pinkish colour in **Figure 3a** is the ERCC4-related helicase is the functional domain helps in replication, recombination and repair.

**Figure 3a:**
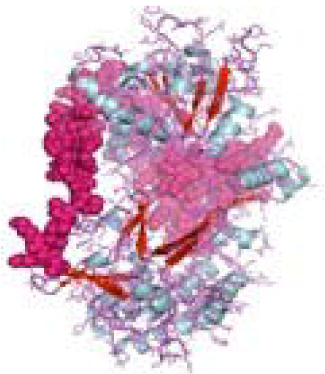
MPH1 ERCC4-related helicase [Replication, recombination and repair] of sheep.

CARD_RIG-I_r2(16-93) is the second repeat of the domain, Caspase activation and recruitment domain found in RIG-I. Similarly, CARD_MDA5_r2(16-93) is the second repeat of Caspase activation and recruitment domain found in MDA5.

CARD domain (6-87) is depicted in **Figure 3b** in the green and pink portion is the Caspase activation and recruitment domain which functions as a protein-protein interaction domain. Lastly, the PRK13766 (678-771) is the Hef nuclease; Provisional domain in a bluish colour in **Figure 3c**.

**Figure 3b:**
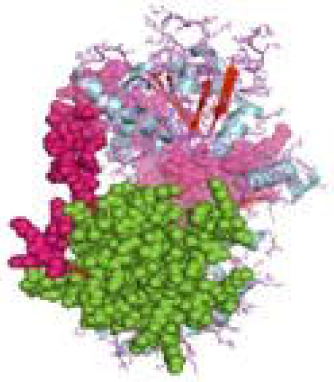
CARD, Caspase activation and recruitment domain: a protein-protein interaction domain of sheep.

**Figure 3c:**
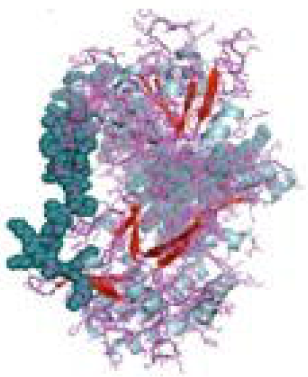
PRK13766, Hef nuclease; Provisional.

### 3.3. Depiction of Protein-protein interaction network and estimation of biological function

The relationship of RIG-I(DDX58 or DExD/H-box helicase 58) with other proteins have been obtained by String analysis **Figure 4**. The related proteins revealed are MAVS (Mitochondrial antiviral signaling protein (520 aa), ISG15 (Ubiquitin-like protein ISG15; Ubiquitin-like protein which has an essential role in the innate immune response to viral infection either via its conjugation to a target protein (ISGylation) or via its action as a free or unconjugated protein. ISGylation has a flow of enzymatic reactions involving E1, E2, and E3 enzymes which catalyze the conjugation of ISG15 to a lysine residue in the target protein exhibits antiviral activity towards both DNA and RNA viruses. The secreted form of ISG15 can-persuade natural killer cell proliferation, induce lymphokine-activated-killer (LAK) activity. TRIM25 (tripartite motif containing 25 (TRIM25), mRNA (631 aa)), MX1 (Interferon-induced GTP-binding protein Mx1; Interferon-induced dynamin-like GTPase shows antiviral activity against rabies virus (RABV), vesicular stomatitis virus (VSV) and murine pneumonia virus (MPV). Isoform 1 but not isoform 2 depict antiviral activity against vesicular stomatitis virus (VSV); Belongs to the TRAFAC class dynamin-like GTP ase superfamily. Dynamin/Fzo/YdjA family (651 aa)), IFIT1 (Interferon-induced protein with tetratricopeptide repeats 1 (474 aa), STAT1 (Signal transducer and activator of transcription 1, 91kDa (STAT1), mRNA (1162 aa)), USP18 (Ubiquitin specific peptidase 18) belongs to the peptidase C19 family, EIF2AK2 (Eukaryotic translation initiation factor 2-alpha kinase 2), MX2 (Interferon induced GTP-binding protein Mx2, Interferon induced dynamin like GTPase with antiviral activity against vesicular stomatitis virus), HERC5 (hect domain and RLD 5).

**Figure 4:**
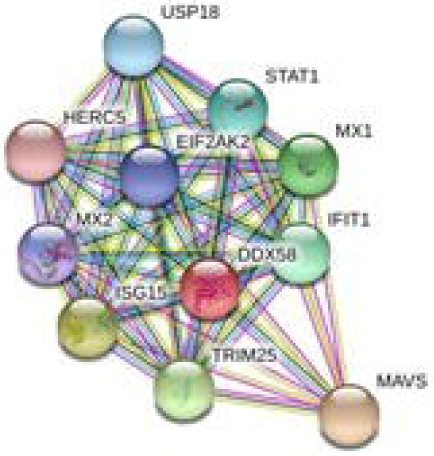
Protein-protein interaction network of RIG1 gene with associated protein by STRING analysis.

### 3.4. In-silico analysis for the detection of the binding site of RIG-I with parasite (*Haemonchus contortus*)—Molecular Docking

Some of the important structural proteins of Haemonchus contortus has been identified and sequences retrieved from gene bank (NCBI). The current section **(Table 5)** discusses the findings obtained from molecular docking of RIG-I with identified domain of H.contortus with the prediction of possible binding site.

**Table 5:**
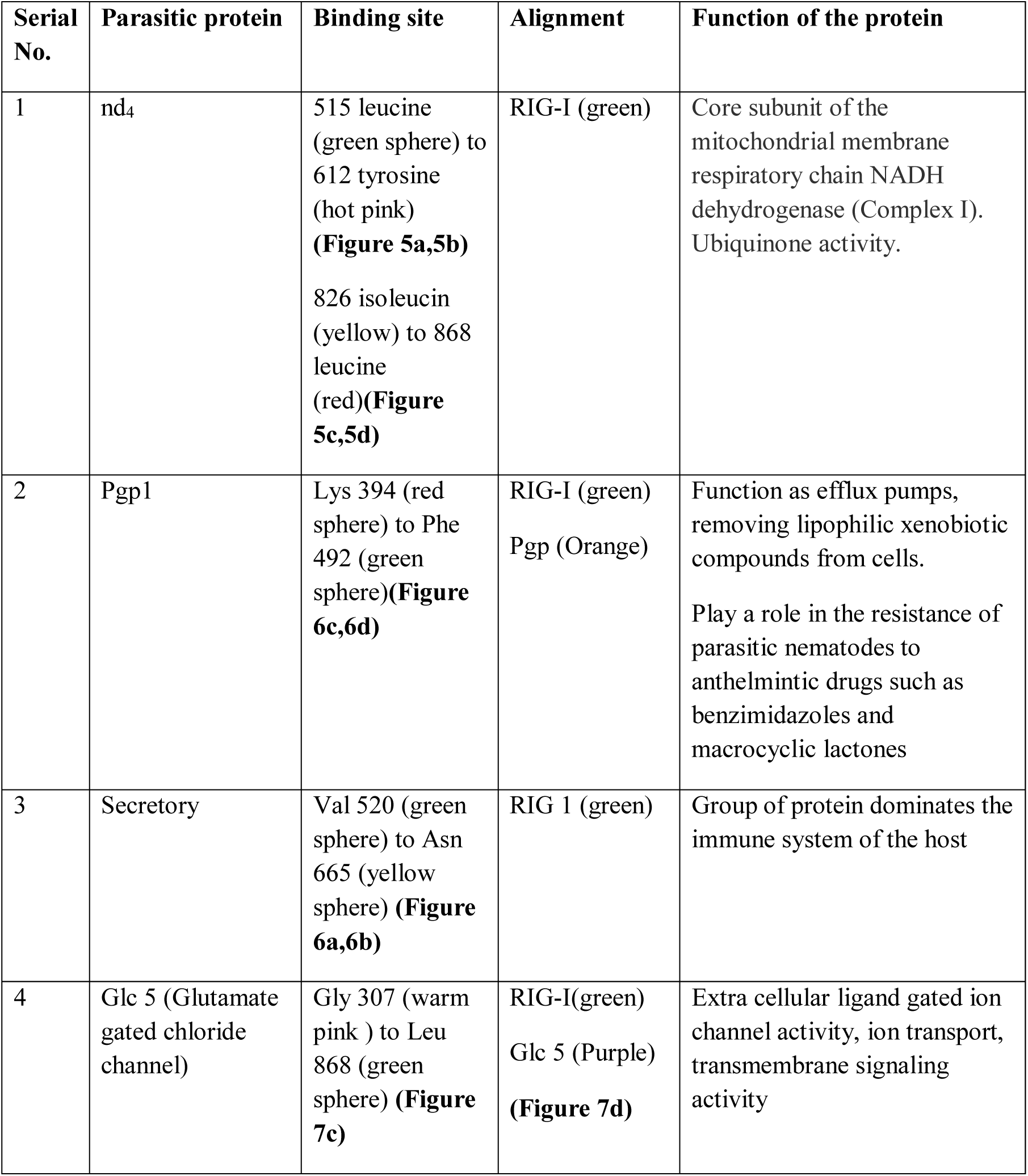
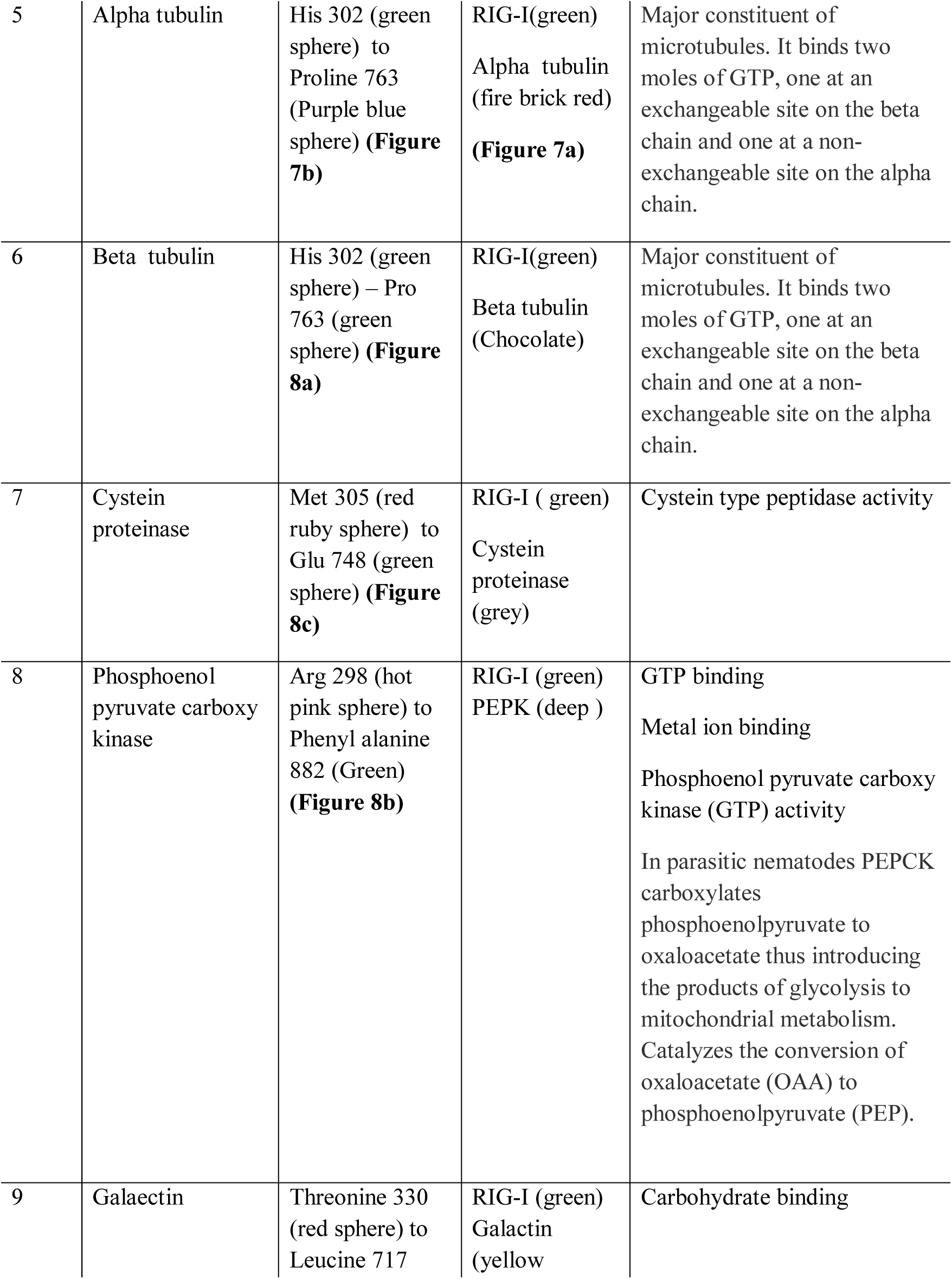
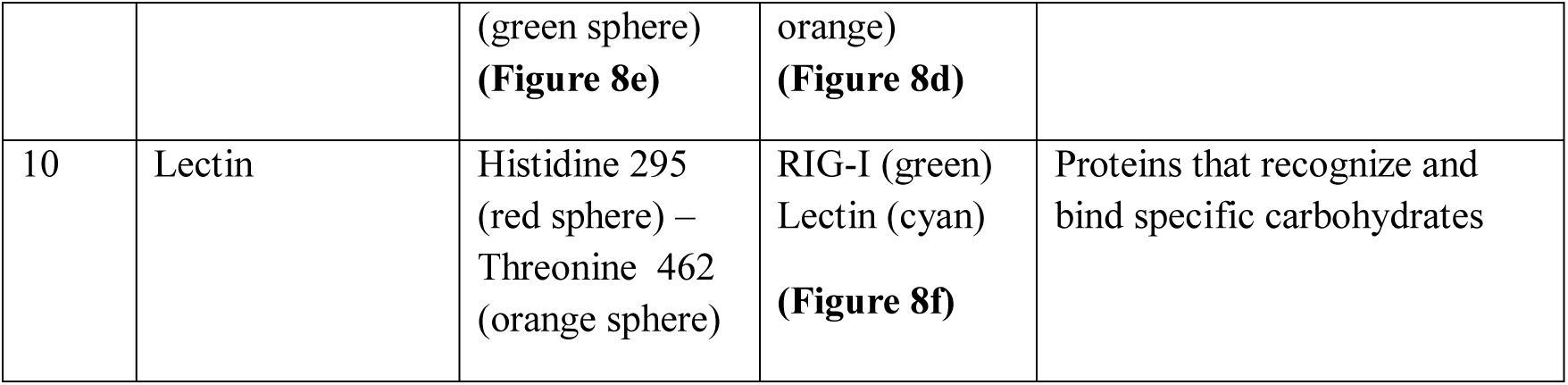
Molecular docking analysis of RIG-I with the protein of *Haemonchus contortus*:

The table shows the binding sites of the Haemonchus protein with the RIG-I protein analysed by molecular docking. Binding of RIG-I of sheep with ND4 domain of *Haemonchus contortus*.

The binding positions of the parasitic protein nd4 has been found in between 515 leucin (green sphere) to 612 tyrosine (hot pink) and 826 isoleucin (yellow) to 868 leucine (red). The pgp1 protein which binds from Lysine 394 (red sphere) to Phenyl alanine 492 (green sphere) possesses ATPase activity. Several secretory parasitic protein which play an important role in dominating the host immune system also has been observed at the position between Valine 520 (green sphere) to Asparagine 665 (yellow sphere). The Glc 5 parasitic protein (Glutamate gated chloride channel) that binds from Glycine 307 (warm pink) to Leucine 868 (green sphere). α-tubulin and β-tubulin being major constituent of microtubules, binds two moles of GTP, one at a non-exchangeable site on the alpha chain and one at an exchangeable site on the beta chain. They bind from the Histidine 302 (green sphere) position to Proline 763 (Purple blue sphere) position. The parasitic protein Cystein proteinase which binds from Methionine 305 (red ruby sphere) position to Glutamate 748 (green sphere). Phosphoenol pyruvate carboxy kinase is yet another important parasitic protein that binds from the position Arginine 298 (hot pink sphere) to Phenyl alanine 882 (Green). Lastly, Galaectin, from postion Threonine 330 (red sphere) to Leucine 717 (green sphere) and Lectin from position Histidine 295 (red sphere) and Threonine 462 (orange sphere) are the proteins that recognizes and binds to specific carbohydrate moieties.

### 3.5. Differential mRNA expression profile of Garole sheep with respect to RIG-I gene in healthy sheep with respect to different body tissues

**Figure 9a** has depicted RIG-I expression profiling for different gut associated lymphoid tissue and lymph node. It shows the expression of mRNA highest in the Abomassum tissue almost five folds than all the other organs. Since we are studying RIGI gene expression in gut associated lymphoid tissue(GALT) in response to Haemonchus contortus infection, it is being observed from Fig 9a, expression is least in lymph node and small intestine. Highest is observed in abomassum followed by rumen, then liver. Caecum reveals comparatively less expression (Fig9a).

**Figure 5a,b,c,d:**
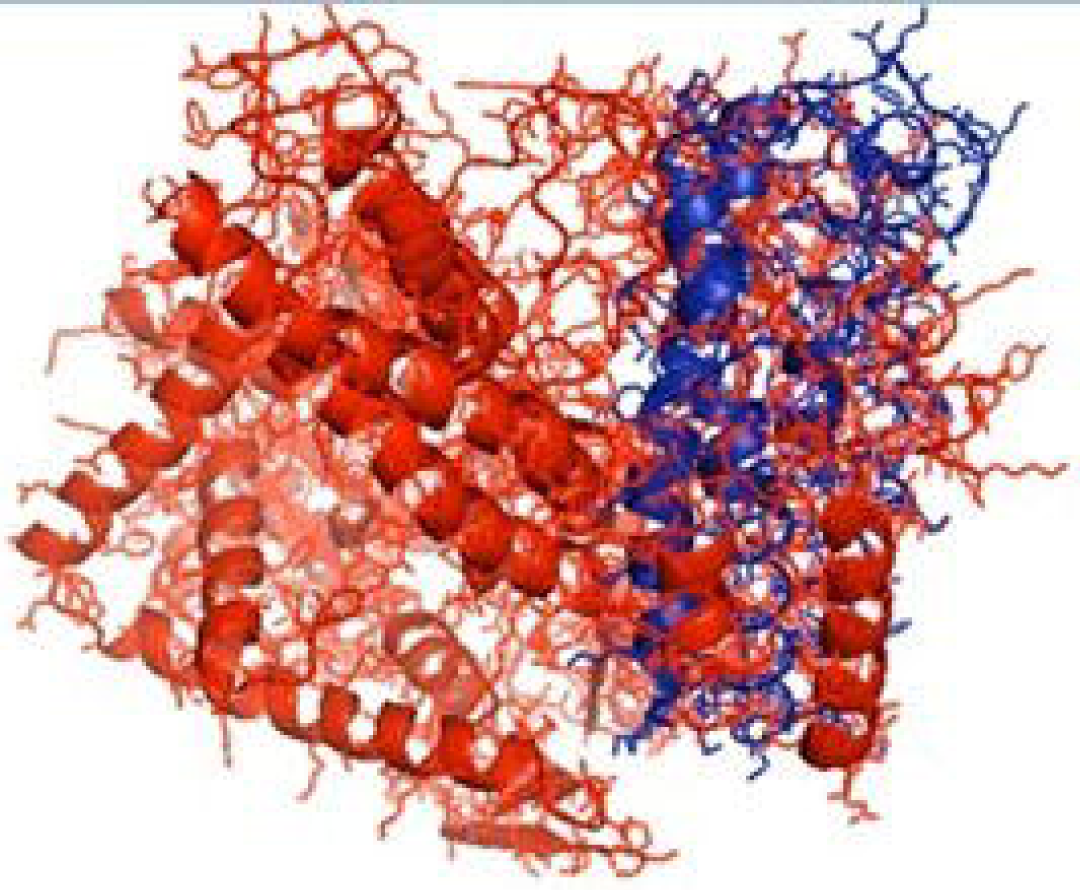
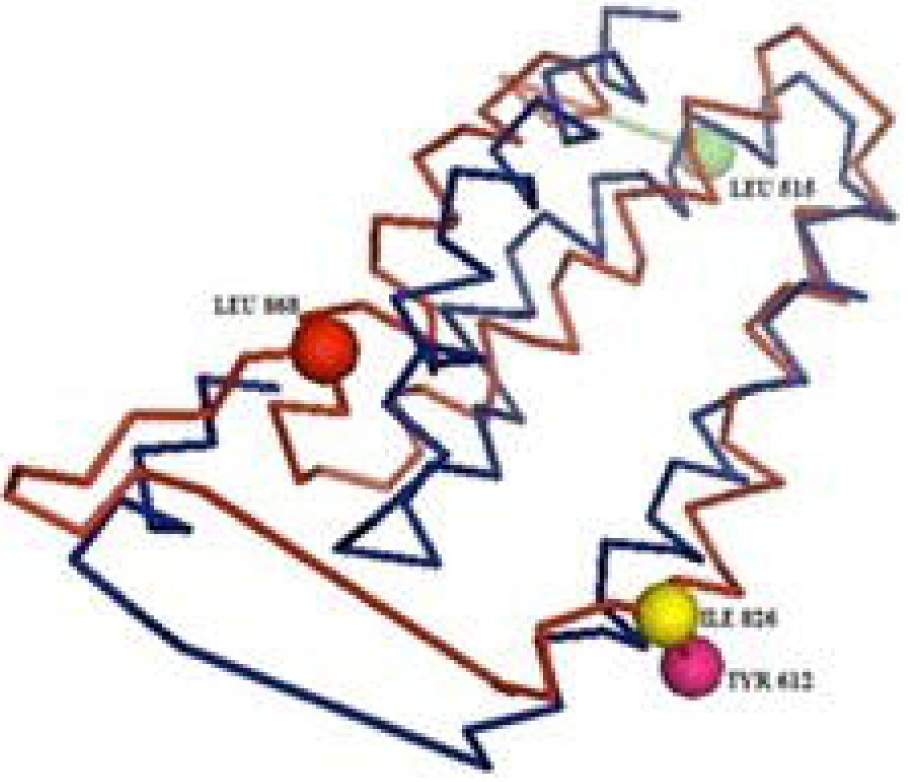
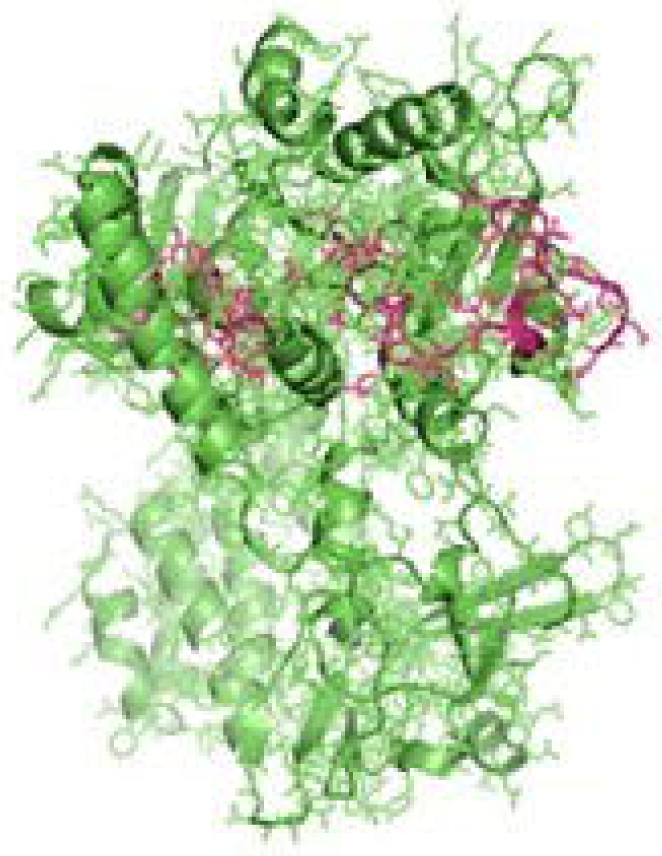
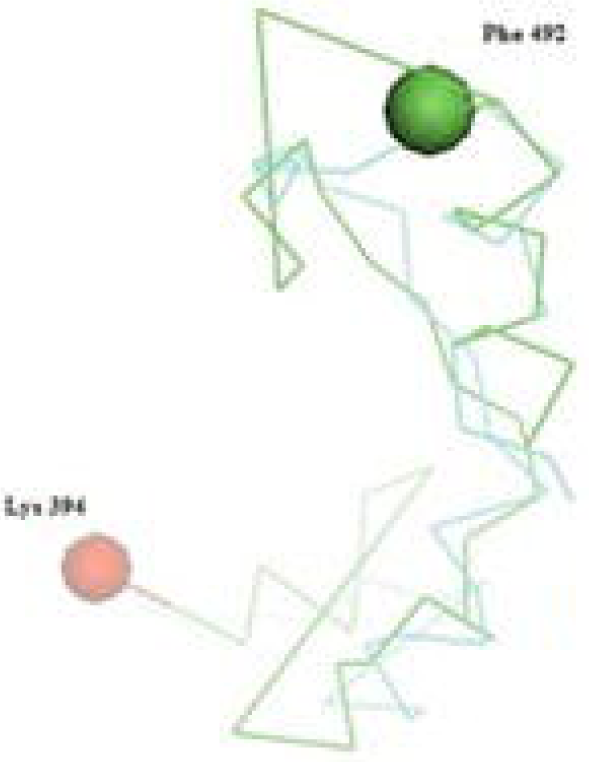
Molecular docking of parasitic protein nd_4_ with ovine RIGI, alignment (left) and binding site (right)

**Figure 6a,b:**
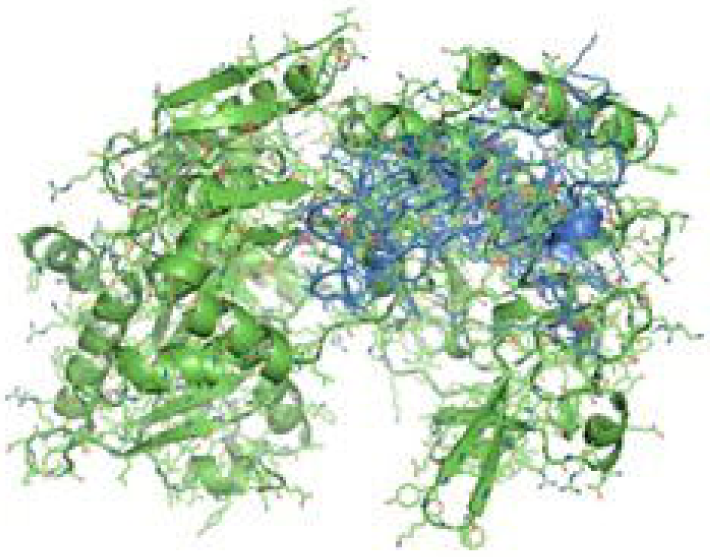
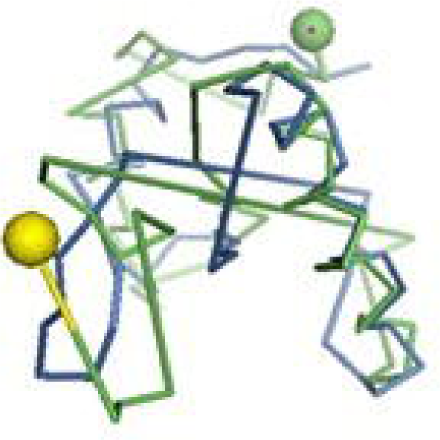
Molecular docking of parasitic protein 15 kDa secretory protein with ovine RIGI, alignment (left) and binding site (right)

**Figure 6c,d:**
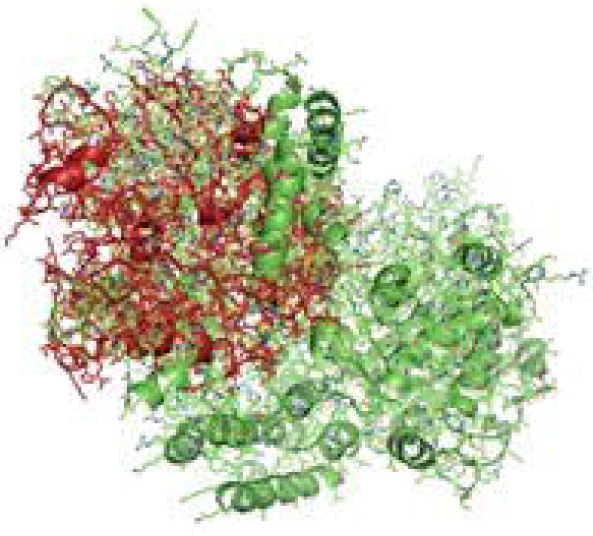
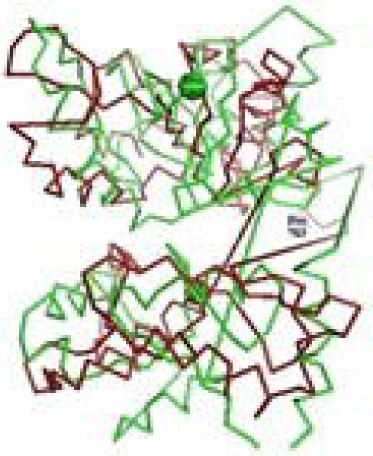
Molecular docking of parasitic protein Pgp1 with ovine RIGI, alignment (left) and binding site (right)

**Figure 7a,b:**
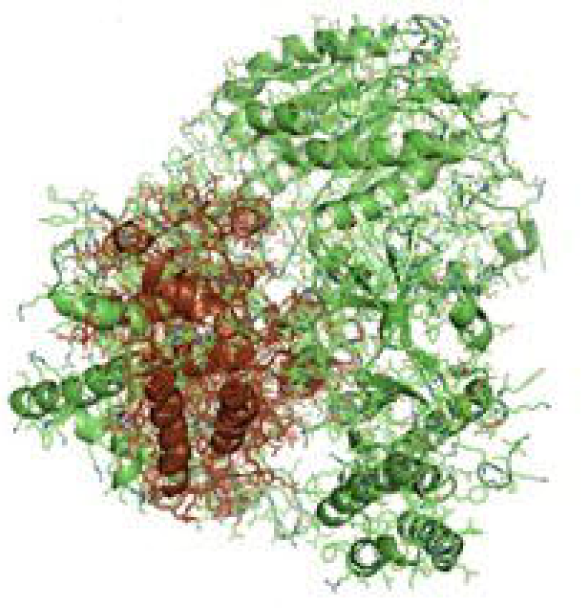
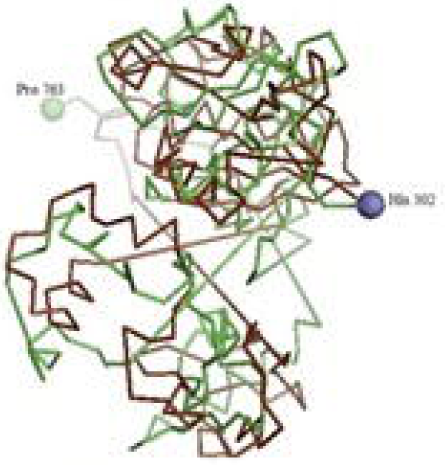
Molecular docking of parasitic protein Alpha tubulin with ovine RIGI, alignment (left) and binding site (right)

**Figure 7 c,d:**
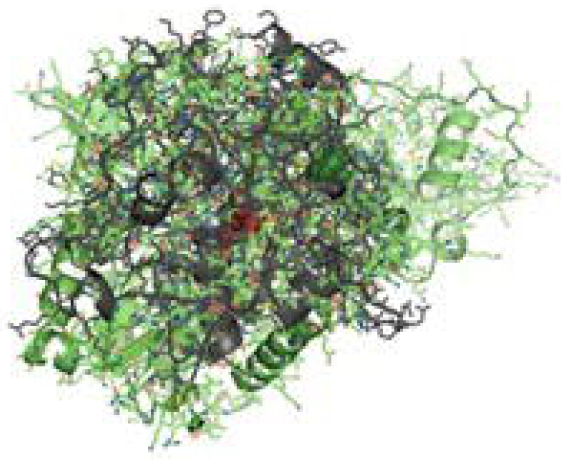
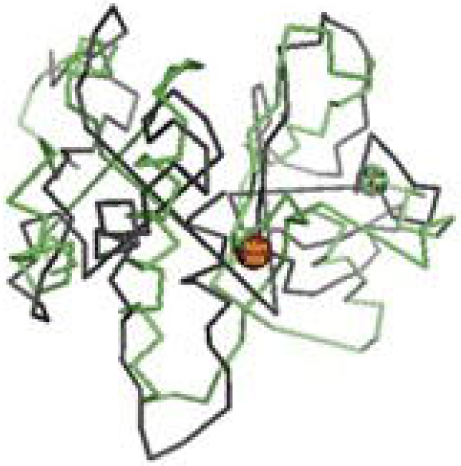
Molecular docking of parasitic protein Glc 5 with ovine RIG-1.

**Figure 8a:**
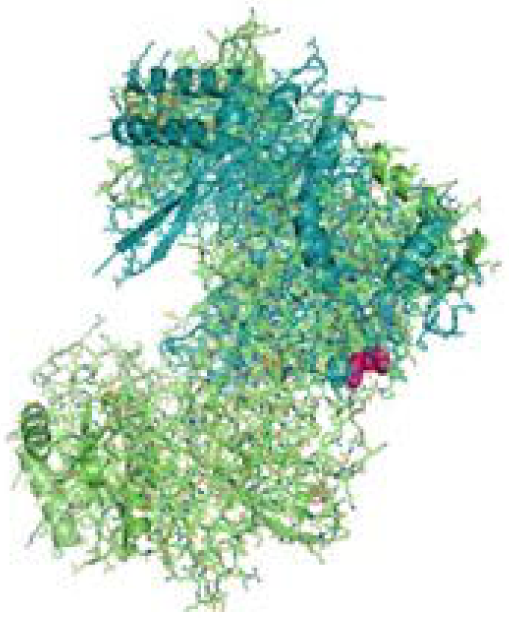
Molecular docking of parasitic protein Beta tubulin with ovine RIGI, alignment (left) and binding site (right)

**Figure 8b:**
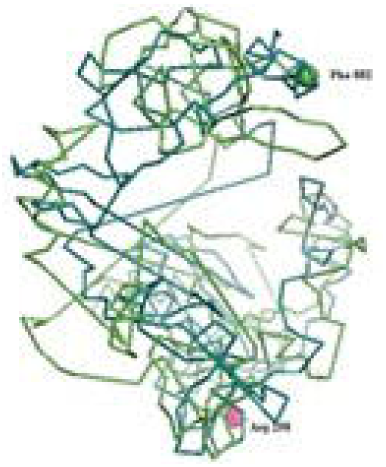
Molecular docking of parasitic protein Phosphoenol pyruvate carboxy kinase with ovine RIGI, alignment (left) and binding site (right)

**Figure 8c:**
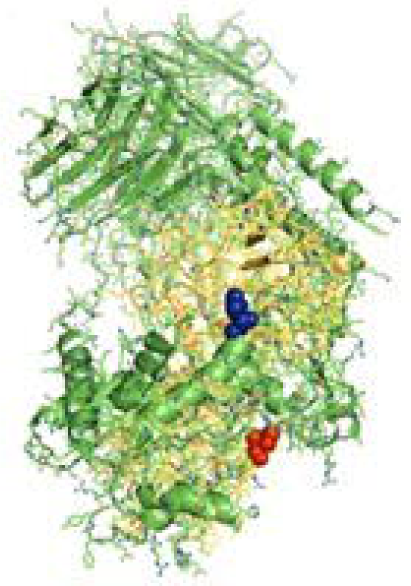
Molecular docking of parasitic protein Cystein proteinase with ovine RIGI, alignment (left) and binding site (right)

**Figure 8d,e:**
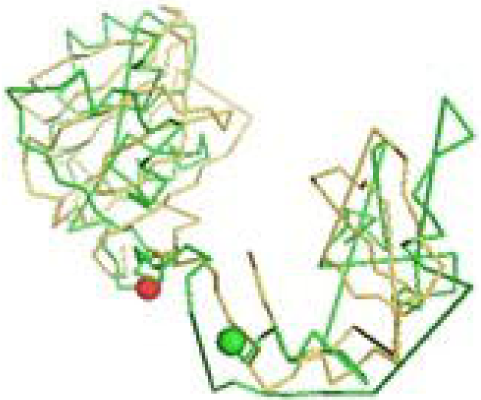
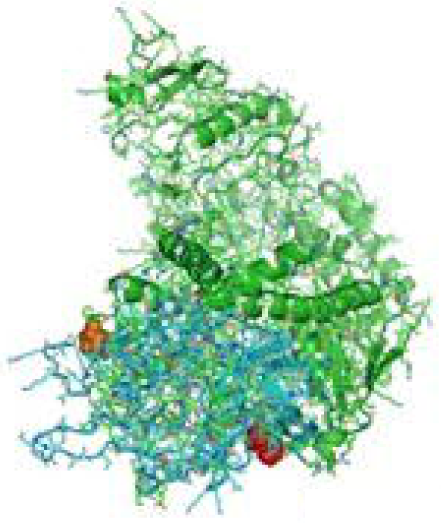
Molecular docking of parasitic protein Galaectin with ovine RIGI, alignment (left) and binding site (right)

**Figure 8f:**
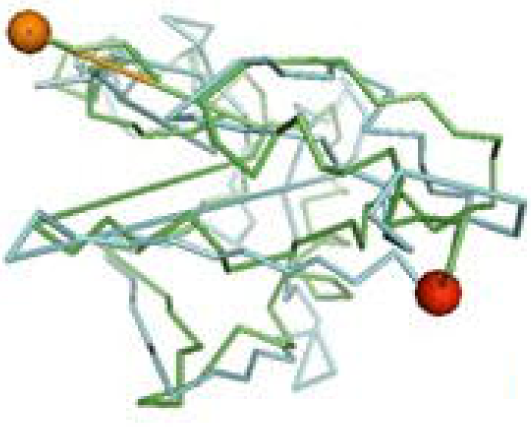
Molecular docking of parasitic protein Lectin with ovine RIGI, alignment (left) and binding site (right)

**Fig 9a:**
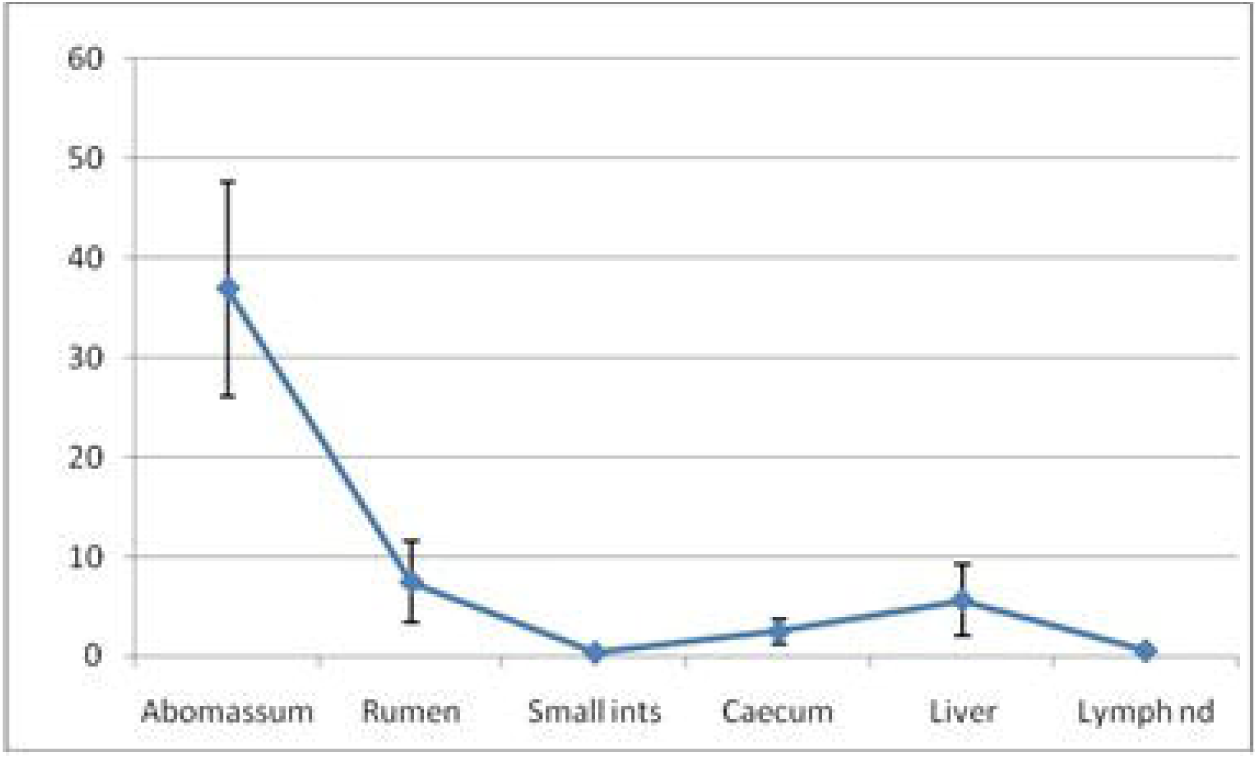
RIGI expression profiling for different gut associated lymphoid tissue and lymph node in sheep.

### 3.6. mRNA expression profile of Garole sheep with respect to RIG-I gene in healthy sheep and those infected with *Haemonchus contortus*

RIG-I expression profiling for healthy and *Haemonchus contortus* infected sheep from abomassum is clearly shown in **Figure 9b**. In the assessment of differential mRNA expression profile of RIG-I genes in Garole sheep with respect to healthy and infected are represented in the figure. RIG-I is better expressed in infected sheep compared to that of healthy sheep of about 2.5 folds.

**Fig 9b:**
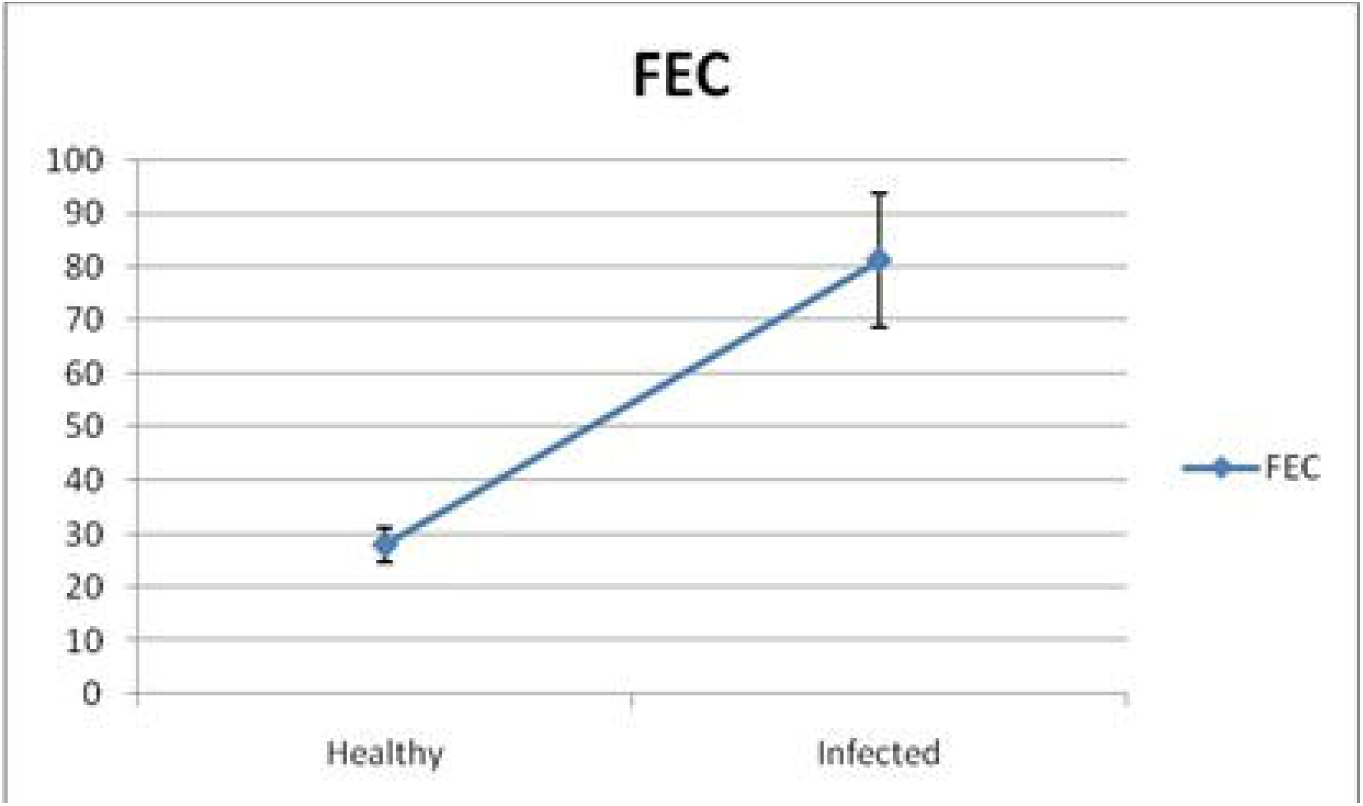
RIGI expression profiling for healthy and Haemonchus contortus infected sheep from abomassum.

### 3.7. Molecular evolution of Sheep with other species with respect to RIG-I gene

The molecular phylogenetic analysis of sheep have been conducted with human, other primates and livestock species as depicted in **Figure 10** with respect to the gene of interest. All the ruminant species are clustered together (goat, cattle, buffalo). Next closely related species are pig and camel. The tree depicts more close genetic relationship of human with sheep, compared to that of lab animals such as rat or mice.

**Figure 10.**
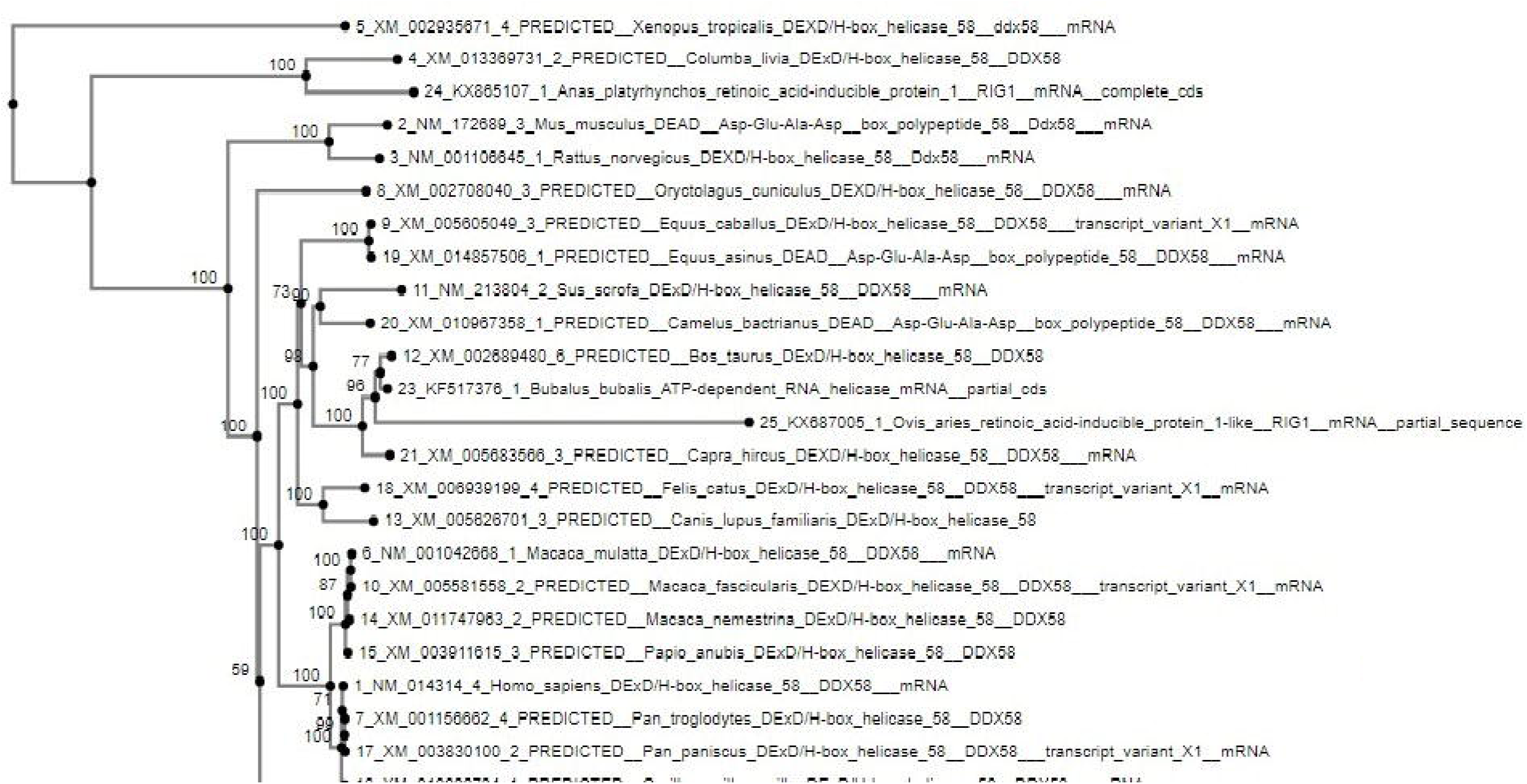
Molecular Phylogenetic analysis of sheep with other species based on RIGI gene.

## 4. Discussion

RIG-I gene which is considered to be a family of the RLR family is known to recognize viral DNA in the cytoplasm. According to previous findings and information, RIG-I was considered to act as an antiviral gene which recognizes double stranded viral DNA and plays an important role in immune regulation. CARD (Caspase recruitment domain) like regions are present in the gene at its N-terminal position which acts as interacting domain with other proteins in which CARD regions are present. It also consists of Repressor domain at the C-terminal position which takes part in RNA binding and the central DExD helicase domain that acts as the ATP binding motif. RIG-I is reported to have a role in immune responses in association with different infectious and non infectious diseases like astherosclerosis, skin psoriasis in which the levels of Interferon gamma and RIG-I is significantly raised in the epidermis of the skin ^**7**^.

Previously, much less is known about the role of RIG-I gene as an anti parasite in sheep, therefore in our experiment we have tried to highlight the fact that RIG-I can be induced in response to parasite infection and can influence the immune regulation against it.

RIG-I expression was observed to be important for gut associated lymphoid tissue (GALT) according to some reports. It is an important component of Musoca associated lymphoid tissue which regulates the immune system of the body and protects the gut from the invasion of infections. Although being a component of immune system, its fragility and permeability makes it vulnerable towards infecting agents. Mostly, parasites invading the animal finds a way through this route in the body. GALT consists of several plasma cells that can produce several antibody which acts as a defence against GI nematodes. Haemonchus, being a fatal blood sucking GI nematode affects the mucus membrane affects the abomasum portion of the sheep^**22**^.

In the present work we have randomly categorized the sheep into two groups as Healthy and disease based on the examination of faecal Egg Count. Sixty sheep were taken into account from which six in number from each group with highest faecal count and lowest faecal count were considered for the statistical analysis. Here, we have shown differential mRNA expression profiling of Abomasum in healthy and Haemonchus infected sheep and observed that differential mRNA expression level for RIG-I was much higher (2.5 folds) in Haemonchus infected sheep in comparison to that of healthy subjects. Abomasum reveals marked increase in lymphoid tissue compared to lymph node. Since lymph node being a secondary lymphoid organ, it plays a role in the adaptive immune system. Due to better expression level and since Haemonchus affects the abomasum severely,we have considered this part to be the most affect part which would show a proper level of gene expression.

This implies the better expression of RIG-I in diseased sheep, and thus involved in immune response. This is the first report of the role of RIG-I in parasitic immunity. To establish the fact and to predict the binding site, we had analyzed with molecular docking for RIG-I with different proteins (mostly surface protein) of Haemonchus contortus. Since this is the first report, mechanism of action of RIG-I is not known against parasitic diseases. Earlier reports indicate that when RIG-I is stimulated with TLR4, it induces its expression in macrophages and gradually depletion of RIG-I causes inhibition of TLR4 induced bacterial phagocytosis^**23**^. Likewise in genes analogous with the early inflammatory response and even those encoding toll-like receptors such as TLR 2, 4 and 9 or in close relationship with free radical production (DUOX1 and NOS2 A) are more profusely expressed in lambs that are resistant to *H.contortus* and *Trichostrongylus colubriformis* infections^**23**^.

Different important domains of RIG-I of *Ovis aries* have been analysed through *in silico* studies. The C-terminal regulatory domain is one of the most functional domain in the gene. It binds with the viral RNA and activation of the RIG-I ATPase by RNA dependent dimerization occurs. RD type is a zinc binding domain that is related to GDP/GTP. Similarly, the molecular kinetics of RIG-I contains two N-terminal CARD Domains and one RNA helicase domain which help in relaying the signal to the downstream signal adaptor mitochondrial antiviral signaling protein or MAVS which leads to the activation of type I IFN responses ^**24**^. MDA5 along with RIG-I results in the binding of RLRs to the MAVS which is a signal adaptor that influences the activation of NFkb gene. Finally LGP2 plays a role in producing antiviral responses against viruses that are recognised by RIG-I.

Similar studies which corresponds to our findings indicate an increase in serum concentrations of IgE, IgA, IgG, TNF-β, IFN-γ, and IL-6, in naturally infected sheep with *Haemonchus* spp. on pastures with two different nutritional conditions ^**22**^ by studying the immune response in sheep by significantly higher peripheral eosinophilia.

The binding sites of the Haemonchus protein with the RIG-I protein are analyzed by molecular docking. The binding positions of the parasitic protein nd4 has been found in between 515 leucine to 612 tyrosine and 826 isoleucin to 868 leucine. Biologically nd4 is the core subunit of the mitochondrial respiratory chain NADH dehydrogenase (Complex I) that is believed to belong to the minimal assembly required for catalysis. Complex I functions in the transfer of electrons from NADH to the respiratory chain. The immediate electron acceptor for the enzyme is assumed to be ubiquinone. Domain analysis for RIG-I had revealed the polypeptide binding site (RD interface) as nucleotide site 522, 525-526, 539-540, 543. Other polypeptide binding site (Helicase domain interface) for RIG-I are 514-515, 518-522. Interestingly it has been observed that the binding site of RIG-I corresponds with the parasitic protein (nd4, alpha tubulin, beta tubulin, cystein proteinase, lectin, galectin, pepk) as revealed through *in silico* studies. The pgp1 protein which binds from Lysine 394 to Phenyl alanine 492 possesses ATPase activity. Pgp1 protein which is also known as a multi drug resistant protein, removing lipophilic xenobiotic compounds from cells functioning as efflux pumps, and also play a role in the resistance of parasitic nematodes to anthelmintic drugs such as benzimidazoles and macrocyclic lactones. Several secretory parasitic protein which play an important role in dominating the host immune system also has been observed at the position between Valine 520 to Asparagine 665. These kind of proteins either depress the host immune system or instigate the system. It has been reported earlier that Haemonchus controtus excretory and secretory proteins (HcESPs) can bind to goat’s peripheral blood mononuclear cells. HcESPs affected the biological functions of the cell, such as cell proliferation, cytokine production, cell migration and nitric oxide production of goat’s PBMCs ^**25**^ .Gradual decrease in Interleukin-4 and Interferon-γ was observed in PBMCs, in contrast there was increase in concentration of IL-10 and IL-17^**26**^. Nitric Oxide production was suppressed at different concentration of HcESPs. Similarly, cell proliferation was also decreased at all concentrations of the parasitic proteins ^**26**^. The Glc 5 parasitic protein (Glutamate gated chloride channel) that binds from Glycine 307 to Leucine 868. These chloride channels performs extra cellular ligand gated ion channel activity, ion transport, transmembrane signaling. *H.controtus* being a highly resistant and polymorphic parasite in livestock is also resistant to Ivermectin. On the other hand *Caenorhabditis elegans* enables to study the effects of polymorphism in resistant parasites. Reports claim that this was tested using glutamate chloride channels that forms candidate resistant genes and an ivermectin drug target ^**27**^.

α-tubulin and β-tubulin being major constituent of microtubules, binds two moles of GTP, one at an exchangeable site on the alpha chain and one at a non-exchangeable site on the beta chain. They bind from the Histidine 302 position to Proline 763 position. They play an important role in Haemonchus controtus by providing resistance to Benzimidazole Antihelmintics ^**28**^. The parasitic protein Cystein proteinase which binds from Methionine 305 position to Glutamate 748. This protein has a peptidase activity and are responsible for breaking down the host haemoglobin. Haemonchus contortus is of the most highly pathogenic parasite of ruminants and cystein proteases acts as one of the prime targets for vaccine against H.controtus. Being an active protease of the excreatory-secretory product of H.controtus, there is strong probability of its involvement in induction of protective immunity ^**29**^. This proteases are also considered as important digestive enzymes in parasitic helminthes.

Phosphoenol pyruvate carboxy kinase is yet another important parasitic protein that binds from the position Arginine 298 to Phenyl alanine 882. This is an important enzyme in helminth parasites since it has GTP binding activity, metal ion binding activity and moreover it catalyses the reverse reaction of formation of Oxaloacetic acid from phosphoenol pyruvate (PEP) rather than in mammals in which the reaction is vice versa. This leads to an important functional difference between the host and the parasite and it can be predicted as a novel target for antihelminthic drugs. These predictions are subjected to experimental validation and discovery of novel chemotherapeutic agents to fight against helminthes, protozoas etc ^**30**^. Lastly, Galaectin, from postion Threonine 330 to Leucine 717 and Lectin from position Histidine 295 and Threonine 462 are the proteins that recognizes and binds to specific carbohydrate moieties.

## 5. Conclusion

RIGI gene of sheep was characterized for the first time. Some important domains for ovine RIGI were identified. So far RIGI gene was studied mostly as an agent conferring antiviral immunity. In this current study, we detected the role of RIGI in conferring immunity against gastrointestinal nematode as Haemonchus contortus. The fact was revealed through differential mRNA expression profile of RIGI in Haemonchus infected in respect to healthy sheep. The fact was later confirmed through in silico studies with molecular docking. Since this is the first report, we had identified the binding site for RIGI with the polypeptide of parasite at 300-750 for most of the protein. Alpha tubulin, Beta tubulin, lectin, galectin and cysteine protease were predicted to be the most promising binding site with ovine RIGI. Molecular phylogeny have detected sufficient genetic similarity of domestic sheep with human, next to primates. Hence sheep may be effective employed as animal model for studying parasitic immunology.

## Acknowledgements

The authors are thankful to Department of Biotechnology, Ministry of Science and Technology, Govt. of India to provide the financial grant vide number No. BT/Bio-CARe/04/10100/2013-14. The authors are thankful to Vice-Chancellor, West Bengal University of Animal and Fishery Sciences and Visva Bharati University for providing lab space and support for carrying out the research work.

## List of Tables

1. List of primers used for amplification of RIG-I gene of Garole sheep as overlapping sequences.
2. List of primers used for QPCR study
3. Mean FEC of the Healthy and Diseased sheep
4. Molecular docking analysis of RIG-I with the protein of *Haemonchus contortus*

## Notes

### Competing Interest Statement

The authors have declared no competing interest.

